# Neural and behavioral similarity-driven tuning curves for manipulable objects

**DOI:** 10.1101/2024.04.09.588661

**Authors:** D. Valério, A. Peres, F. Bergström, P. Seidel, J. Almeida

## Abstract

In our daily activities, we encounter numerous objects that we successfully distinguish and recognize within a fraction of a second. This holds for coarse distinctions (e.g., cat vs. hammer) but also for more challenging within-category distinctions that require fine-grain analysis (e.g., cat vs. dog). The efficiency of this recognition depends on how the brain organizes object-related information. While several attempts have focused on unravelling large-scale organization principles, research on within-category organization of knowledge is rather limited. Here, we explored the fine-grain organization of object knowledge and investigated whether manipulable objects are organized and represented in terms of their similarity. To accomplish this, different groups of individuals participated in a behavioral and fMRI release from adaptation experiment. Adaptation was induced by presenting different exemplars of a particular object, and release from adaptation was elicited by the presentation of a deviant object. The relationship between adaptation and deviant objects was manipulated into four levels of similarity, measured by feature overlap between these objects. Our findings revealed that increasing object similarity provoked progressively slower reaction times and progressively weaker fMRI release from adaptation. Specifically, we identified similarity-driven tuning curves for the release from adaptation in the medial fusiform, collateral sulcus, parahippocampal gyri, lingual gyri, lateral occipital complex, and occipito-parietal cortex. These results suggest that the processing and representation of objects in the brain and our ability to perform fine discriminations between objects reflect real-world object similarity in a relatively parametric manner.

## Background

One of the most challenging questions in cognitive neuroscience centers on understanding how we efficiently recognize and distinguish objects. To unravel this ability, we must delve into how information is represented and organized in the brain (e.g., Almeida et al., 2023). A widely accepted hypothesis suggests that information might be organized based on similarity in external world dimensions (Edelman, 1998). Examples of this organizational principle abound in sensory-motor cortices – for instance, the visual cortex is organized according to the location of the stimuli in the visual field (i.e., retinotopic maps; Sereno et al., 1995), and the auditory cortex according to the tone of an auditory stimulus (i.e., tonotopic maps; Formisano et al., 2003). In line with this, several studies showed that similarity matters for conceptual representation in the brain – these studies focused on a coarse, between-category, representational level (e.g., Kriegeskorte et al., 2008). Whether this organization principle extends to finer-grained within-category object distinctions remains unclear. Here, we investigated whether within-category object similarity shapes the organization of neural and cognitive object representations.

Exploring how object knowledge is represented in the brain has led to the proposal of several hypotheses on the organization of information within the ventral and lateral temporal cortex (Kanwisher et al., 1997; Caramazza and Shelton, 1998; Chao et al., 1999; Haxby et al., 2001; Levy et al., 2001; Epstein, 2008; Kriegeskorte et al., 2008; Almeida et al., 2013; Konkle and Caramazza, 2013; Sha et al., 2015; Downing and Peelen, 2016; Mahon and Almeida, 2024 for a review see Grill-Spector and Weiner, 2014). For instance, it has been suggested that object knowledge is organized according to coarse domain distinctions, such as animate or inanimate (Caramazza and Shelton, 1998; Kriegeskorte et al., 2008), or more specific categories, such as faces, bodies, animals, manipulable objects, places, among others (Kanwisher et al., 1997; Chao et al., 1999; Epstein, 2008; Almeida et al., 2013; Downing and Peelen, 2016); or that the organization of object knowledge follows specific dimensions such as visual form (Haxby et al., 2001), eccentricity preferences (Levy et al., 2001; Arcaro et al., 2019), real-world object size (Konkle and Caramazza, 2013), animacy (Sha et al., 2015), or is organized according to multidimensional arrangements (Huth et al., 2012; Hebart et al., 2020; Fernandino et al., 2022; Almeida et al., 2023a). Importantly, several of these studies suggested that object similarity in the real world drives the organization of conceptual information in the brain (e.g., Kriegeskorte et al., 2008; Huth et al., 2012; Carlson et al., 2014; Bracci and Op de Beeck, 2016; King et al., 2019; Fernandino et al., 2022). Moreover, most, if not all, of these studies have tested the role of similarity at a coarse, between-category level.

The number of studies evaluating the within-category organization of object knowledge is considerably lower – that is, studies investigating how information is represented among basic level members of a single domain (e.g., within the domain of manipulable objects, how is information represented among the different basic level members such as a knife and a pair of scissors; (Gotts et al., 2011; Connolly et al., 2012; Bruffaerts et al., 2013; Bracci et al., 2017; Almeida et al., 2023a, 2023b; Xu et al., 2023). This is at odds with the fact that the hardest and perhaps most common everyday decisions we tend to make (e.g., is the object in front of us a knife or a fork) typically rely on finer-grained, within-category representations. Moreover, brain-damaged patients with category-specific deficits do not struggle with between-category distinctions (fork vs. dog) but rather with within-category distinctions (fork vs. spoon; Caramazza and Shelton, 1998; Moss and Tyler, 2000). This strongly suggests that unravelling finer-grained levels of representation and, specifically, focusing on whether object similarity is central for within-category object processing is an essential step in understanding object recognition.

Here, we focused on the category of manipulable objects to test whether the organization of within-category representations conforms to object similarity such that progressive differences in similarity are reflected in behavioral responses and neural representations. To achieve this, we conducted behavioral and functional magnetic resonance imaging (fMRI) experiments using a release from adaptation paradigm, where we parametrically manipulated the similarity between the adaptation and deviant object stimuli. We used this paradigm because it allows the extraction of behavioral and neural similarity-driven tuning curves for manipulable objects. In the behavioral experiment, we measured how quickly and accurately participants detected the deviant object. We expected that the more similar the adaptation and deviant objects were to each other, the harder it would be to differentiate them and the longer reaction times and higher error rates would be. In the fMRI experiment, we measured neural responses to the deviants. We anticipated that some regions would exhibit a differential progressive release (after adaptation) as a function of the similarity between adaptation and deviant objects. Particularly, medial aspects of the ventral temporal cortex, associated with surface and material properties processing (Cant et al., 2009; Cavina-Pratesi et al., 2010a, 2010b; Sun et al., 2015), the Lateral Occipital Complex (LOC) involved in processing complex shapes (Grill-Spector et al., 1999), the middle temporal gyrus, responsible for processing functional goals of objects (Lingnau and Downing, 2015; Wurm and Lingnau, 2015), and dorsal occipital cortex and occipito-parietal regions, involved in 3D processing and object manipulation (Jeannerod et al., 1994; Cavina-Pratesi et al., 2007; Monaco et al., 2011; Freud et al., 2016), should show this progressive release from adaptation. As predicted, progressive increases in similarity between adaptation and deviant objects led to progressively and significantly slower reaction times and lower recovery of the BOLD signal in the ventral temporal and occipito-parietal regions.

## Experimental Design

For Experiments 1 (a and b) and 2, we selected a set of manipulable objects previously used in our laboratory (Almeida et al., 2023a, 2023b; Valério et al., 2023). We chose to investigate manipulable objects because 1) these items represent everyday manmade objects that we constantly perceive and interact with; 2) these items encompass different sets of associated information, such as their function, the movements associated with their manipulation, specific structural features (e.g., a handle or rounded shape), and particular contexts in which we use them; 3) this category of objects can be independently impaired (or spared) in patients with brain lesions in the context of spared (or impaired) recognition of items from other domains of knowledge (e.g., animals or faces; for a review see Capitani et al., 2003); and 4) this category is well-suited for using a release from adaptation paradigm because the same object can be rendered in a wide diversity of materials, colors, and shapes (e.g., the various types of glasses). This allows inducing adaptation to an object across different mid-level properties.

To compute between-object similarity, we calculated the cosine similarity formula based on object features and their production frequencies as obtained in a previous study (Valério et al., 2023). In that study, we asked 130 participants to freely generate features for 80 manipulable objects (see Figure 1A; Valério et al., 2023). Note that cosine similarity ranges from 0 to 1, and values closer to 1 indicate high object similarity. Based on these values, we selected pairs of objects according to four similarity levels and binned them together (see Figure 1A right). Further details of each experimental design are provided below and can be seen in Figure 1 (B, C, and D).

**Figure 1.**
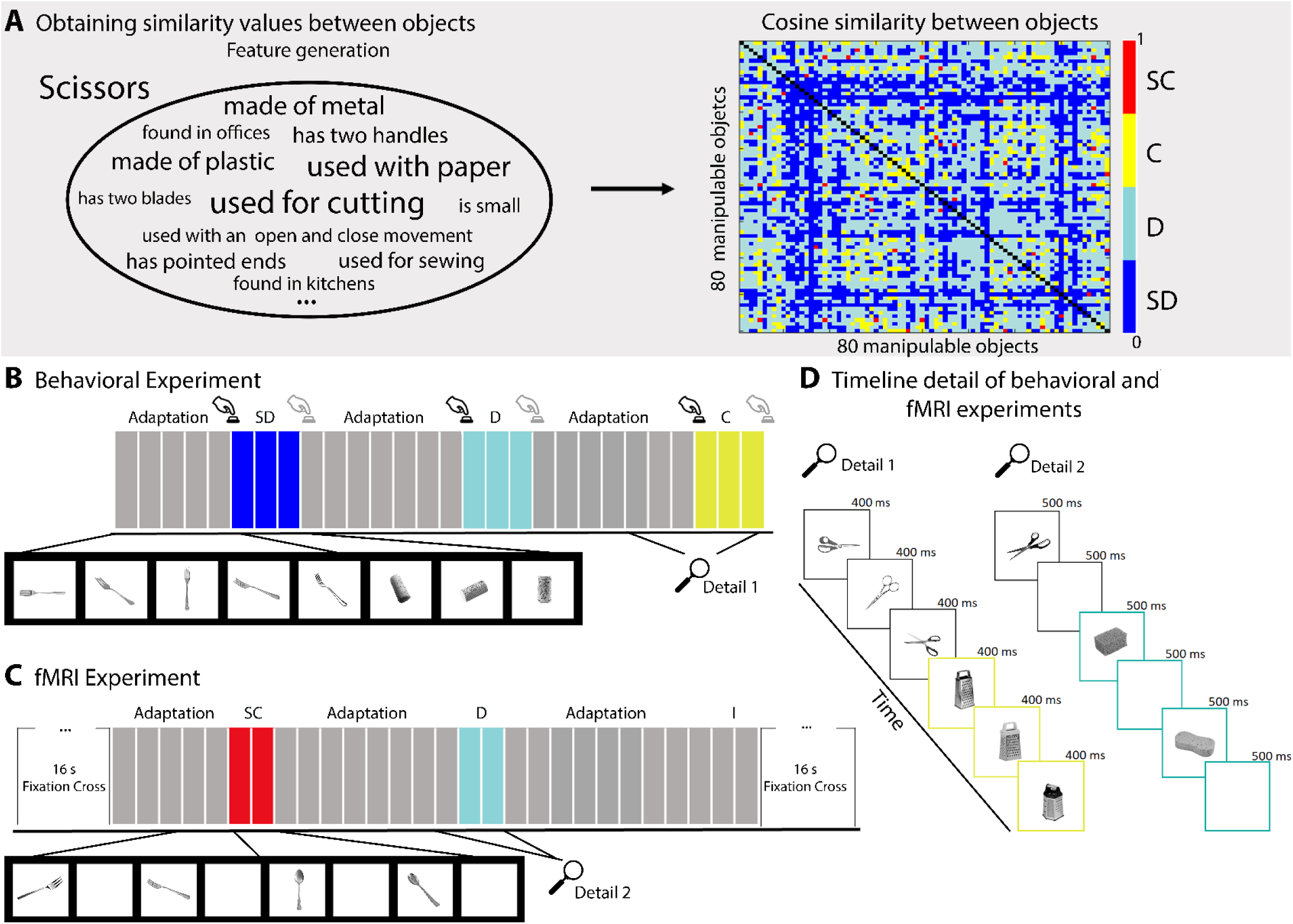
Experimental Design. **A)** In the gray shadow are the data obtained in a previous study (Valério et al., 2023), illustrating the features generated for pairs of scissors. Using these features, we computed cosine similarity scores among 80 objects, considering their production frequency (the font size in the word cloud represents the production frequency values). In the upper right corner, we present an 80 by 80 matrix of cosine similarity values between an object and all the other 79 objects, color-coded according to their similarity. From this matrix, we selected twenty objects – i.e., Experiment 1a and 1b - that contained object pairs fitting into four levels of similarity: those that were very similar to each other in red– i.e., the Super Close bin (SC); those that were similar to each other in yellow – i.e., the Close bin; those that were distant to each other in terms of similarity in light blue – i.e., the Distant bin (D); and those that were very dissimilar to each other in dark blue – i.e., the Super Distant bin (SD). **B)** Design of the behavioral experiment. We presented adaptation sequences (i.e., different exemplars of the same object such as a fork) followed by a deviant object. Participants were instructed to press a button whenever there was a change in the object (e.g., from a fork to a cork) but not when the same object was presented under a different perspective or with a different exemplar of the same object (e.g., two different forks). We recorded responses when the object changed from an adaptation sequence to a deviant, but changes from a deviant to a new adaptation sequence were not analyzed (indicated by a grey button press icon). The number of adaptors in an adaptation sequence varied pseudo-randomly between five to nine. Each image was presented for 400 ms, and each deviant type was presented three times in a row (i.e., different exemplars of the same object) before another adaptation sequence began. **C)** Design of the fMRI experiment. Each run started and finished with 16s of a screen-centered fixation cross. For each trial, each image was presented for 500 ms followed by 500 ms of a blank screen. The number of adaptors in an adaptation sequence varied pseudo-randomly between five to nine, and each deviant appeared twice. Participants were instructed only to look at the pictures. In both experiments, besides the four experimental manipulations of similarity, we included an Identity condition (I), where the adaptation sequence was immediately followed by a different exemplar of the same object. **D)** Two details of the timelines of the behavioral (B) and the fMRI (C) experiments.

### Experiment 1 – Behavioral Experiments

For replicability purposes, we ran this experiment twice (Experiment 1a and Experiment 1b), using the same procedures but using different object pairs and two independent groups of participants.

Methods Participants:

In Experiment 1a, 22 healthy volunteers (16 females; Mean (M) age = 23, Standard Deviation (SD) age = 6.9, range 18 – 47; one left-handed), and in Experiment 1b, 20 volunteers (13 females, M age = 19.9, SD age = 3.4, range 18 - 34; all right-handed) participated. All participants were naïve about the goal and provided written informed consent. All procedures followed ethical guidelines, and the experiments were approved by the Ethics Committee of the Faculty of Psychology and Educational Sciences of the University of Coimbra in accordance with the Declaration of Helsinki.

#### Procedures

Experiments 1a and 1b followed the same procedure and comprised 500 trials each. Each trial consisted of five to nine exemplars of the same object type (e.g., scissors) that appeared in different perspectives and exemplars (adaptation objects), followed by three exemplars of a deviant object (e.g., graters). There were 100 trials for each adaptation length (i.e., five, six, seven, eight, or nine adaptation objects). For each experiment (1a and 1b), we chose 10 adaptation objects paired with five types of deviants according to their similarity. The five deviant conditions consisted of objects that were very similar to the adaptation object (SC condition), similar to the adaptation object (C condition), weakly similar to the adaptation object (D condition), very weakly similar to the adaptation object (SD condition), or different exemplars of the same object (Identity condition). For example, take scissors as an adapting stimulus, paired with five deviants: knife represented the SC condition, grater represented the C condition, sponge represented the D condition, match represented the SD condition, and different exemplars of scissors were the Identity condition. Note that the Identity condition was not of interest in this experiment but was still presented here in order to maintain maximum similarity to the design of Experiment 2. Each deviant condition appeared 100 times. Different pairs of objects were used in Experiment 1 (a and b) for replication purposes with a completely different set of items (and participants) - see Table S1 of the supplementary materials for all pairs used.

All stimuli were gray scaled images of objects. Each image appeared on screen for 400 milliseconds (ms), with a screen refresh rate of 60 Hz. Object size was 400 by 400 pixels, roughly subtending 9.2 degrees of the visual angle. Participants were instructed to perform a one-back task, where they had to detect when the object changed – that is, participants should press a button, as fast as possible, when the displayed image was of a different object than the immediately preceding one. Responses were collected with a Cedrus button box RB-740, and participants used their dominant hand to respond. We measured accuracy and reaction times, and the experiment lasted for 30 min. Participants were not aware that we only collected responses when the object changed from an adaptor to a deviant (as represented by the black button press icon in the Figure 1B), and not vice versa. We used Matlab R2019b and “A Simple Framework” (Schwarzbach, 2011) to present the stimuli.

#### Statistical analysis

We considered trials with reaction times between 100 ms and 800 ms as valid. We chose 800 ms as upper limit because, in that case, participants saw two images in a row without pressing the button. Reaction times faster than 100 ms or slower than 800 ms were considered misses. Overall, 2.1% and 3.2% of the total trials in Experiment 1a and 1b were considered misses (either because the reaction times were below or above the cutoffs described above, or because no responses were obtained), respectively. Reaction times and accuracy values were entered in two separate within-subject one-way ANOVAs, with deviant type (the four deviant types of interest) as a factor. Simple effects between deviant type were conducted for each ANOVA if the factor “deviant type” was significant. All comparisons were Bonferroni corrected and conducted in SPSS (version 22, IBM Corp., Armonk, N.Y., USA, 2013).

## Results

For Experiment 1a, reaction times varied significantly as a function of the type of deviant (*F*(3,63) = 52.08; *p* < 0.0001; and partial η² = 0.71). Planned comparisons showed that participants were slower in the SC (Mean (M) = 448.5; Standard Error Mean (SEM) = 6.4) condition than in the other conditions (C (*t*(21) = 6.83, *p* < 0.0001), D (*t*(21) = 10.99, *p* < 0.0001, and SD (*t*(21) = 12.18, *p* < 0.0001)); there were no reaction time differences between C (M = 425.6; SEM = 6.6) and D (M = 424.6; SEM = 6.8; t(21) = 0.32, *p* = 0.75), but participants were significantly faster in SD (M = 416.7; SEM = 6.7) than in C (*t*(21) = 3.60, *p* = 0.002) and D (*t*(21) = 4.02, *p* = 0.001) conditions. We additionally analyzed the percentage of missing trials (i.e., when participants did not press the button or did so outside of the 100 to 800 ms range). We found that performance varied significantly as a function of the deviant type (*F*(3,63) = 4.91; *p* = 0.004; and partial η² = 0.19). Participants missed more object changes when deviants were part of the SC bin (M = 11.5; SEM = 0.9) than the C (M = 8.4; SEM = 0.9; *t*(21) = 3.05, *p* = 0.006) and SD (M = 8.3; SEM = 0.7; *t*(21) = 2.96, *p* = 0.007) bins. However, we did not find statistical differences between SC and D (t(21) = −1.86, *p* = 0.08), C and D (t(21) = −1.55, *p* = 0.14), C and SD (t(21) = 0.10, *p* = 0.92), or D and SD (t(21) = 1.64, *p* = 0.12).

Importantly, in Experiment 1b we replicated our results with a completely new set of object pairs and an independent group of participants. Performance in reaction time (*F*(3,57) = 29.98; *p* < 0.0001; and partial η² = 0.61) and accuracy (*F*(3,57) = 25.85; *p* < 0.0001; and partial η² = 0.58)) varied significantly by deviant type. Participants were slower in SC (M = 452.3; SEM = 6.6) in comparison to other conditions (C (*t*(19) = 5.44, *p* < 0.0001); D (*t*(19) = 5.22, *p* < 0.0001); and SD (*t*(19) = 8.28, *p* < 0.0001)); there were no differences between C (M = 428.3; SEM = 7.8) and D (M = 433.11; SEM = 7.8; t(19) = - 1.7, *p* = 0.11) conditions; and participants were significantly faster in SD (M = 422.1; SEM = 7.1 in comparison to D (*t*(19) = 4.21, *p* < 0.0001) and SC conditions. Moreover, participants were less accurate in SC (M = 19.70; SEM = 1.8) than C (M = 9.7; SEM = 0.9; *t*(19) = 6.39, *p* < 0.0001), D (M = 10.7; SEM = 1.1; *t*(19) = 5.35, *p* < 0.0001) and SD (M = 8.8; SEM = 0.9; *t*(19) = 5.28, *p* < 0.0001) conditions. However, we did not find statistical differences between C and D (t(19) = −1.16, *p* = 0.26), C and SD (t(19) = 0.93, *p* = 0.36), and D and SD (t(19) = 2.29, *p* = 0.03). All results presented in Table 1 and Figure 2 were Bonferroni corrected for multiple comparisons. See Figure S1 of the supplementary material for the reaction times per type of deviant and participant for Experiment 1 (a and b).

**Figure 2:**
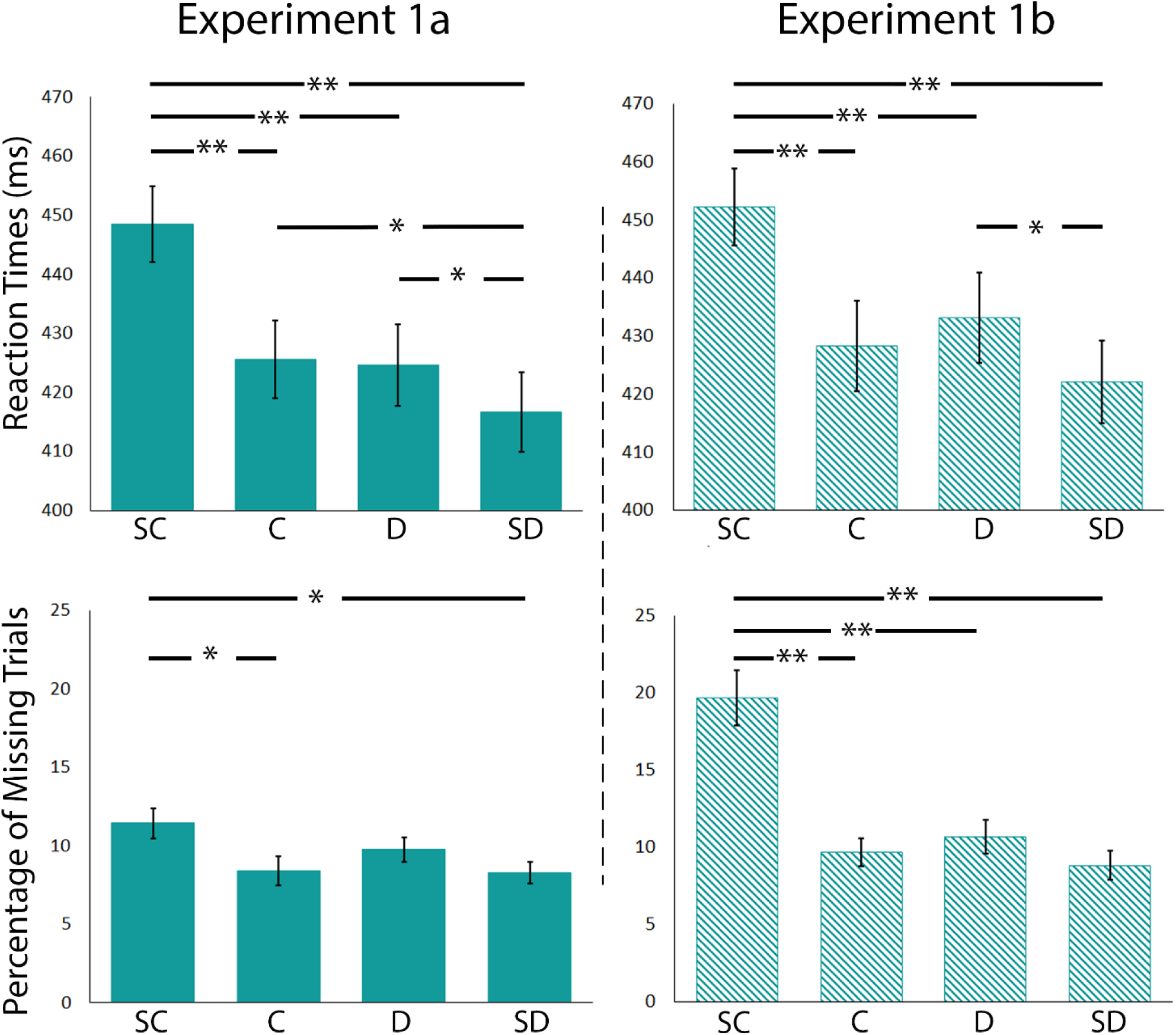
Reaction Times and Percentage of Missing Trials for Experiment 1a and 1b. Here we present reaction times and percentage of missing trials as a function of the similarity between the adaptation and deviant objects. Error bars correspond to the standard error of the mean. SC – Super Close; C – Close; D- Distant; SD – Super Distant *\*\* p* < 0.001, Bonferroni corrected; * *p* < 0.05, Bonferroni corrected

**Table 1:** Reaction Times and Percentage of Missing trials in the Behavioral Experiments.

|  |  | SC |  | C |  | D |  | SD |  |
| --- | --- | --- | --- | --- | --- | --- | --- | --- | --- |
|  |  | Mean | SEM | Mean | SEM | Mean | SEM | Mean | SEM |
| Experiment 1a | RTs (ms) | 448.5 | 6.4 | 425.6 | 6.6 | 424.6 | 6.8 | 416.7 | 6.7 |
|  | Missings (%) | 11.5 | 0.9 | 8.4 | 0.9 | 9.8 | 0.8 | 8.3 | 0.7 |
| Experiment 1b | RTs (ms) | 452.3 | 6.6 | 428.3 | 7.8 | 433.1 | 7.8 | 422.1 | 7.1 |
|  | Missings (%) | 19.7 | 1.8 | 9.7 | 0.9 | 10.7 | 1.1 | 8.8 | 0.9 |

## Discussion

Overall, as anticipated, Experiment 1 (a and b) revealed that participants are slower and less accurate in detecting object changes when two objects share more features. The results demonstrated a relatively graded and progressive effect of object similarity on object detection reaction times and response accuracy - the further a deviant object is from the adaptation object in terms of object similarity, the easier and faster it is to detect an object change. However, this graded effect of object similarity presented a plateau in mid-similarity values – that is, similarity effects were more pronounced for the SC deviants, followed by both the C and D deviants, and were weaker for the SD deviants. This will be discussed further in the General Discussion section.

Our behavioral findings suggest that object similarity figures critically as a principle of organization of the representations within our object conceptual space and do so maintaining a graded nature imposing different levels of object similarity. However, the question remains: how does object similarity relate to the neuronal tuning functions of object-preferring regions?

### Experiment 2 - fMRI Experiment

To investigate whether object similarity influences the neural organization of object knowledge at a fine-grain representational level, we modified the behavioral experiment for an fMRI setup. Specifically, we again used a release from adaptation paradigm, leveraging the well-known fact that neural signals in regions responsible for a particular type of information adapt after continuous and repeated presentation of that information (Grill-Spector and Malach, 2001; Grill-Spector et al., 2006).

In Experiment 2, we expected that brain areas responsible for processing and storing object knowledge would exhibit two neural phenomena: 1) adaptation (a decrease in the BOLD response) to the repeated presentation of exemplars of the same object type (adaptation objects), and 2) recovery from this adaptation (an increase in BOLD response) when presented with a deviant object. It is important to note that these two analyses are statistically independent, as they are conducted on nonoverlapping sets of stimuli and timings of the experiment. Furthermore, what we are interested in is the degree of recovery from adaptation and whether the magnitude of this recovery is a (progressive) function of the similarity between the adaptation and deviant objects – that is, this approach focuses on the voxels that present adaptation and how they recover the BOLD response based on object similarity. What we expect then is that if the neural representation of the adaptation object and the deviant object are extremely similar within a voxel, as expected, for identical or highly related objects with numerous shared features (i.e., highly similar objects), then the recovered response should be relatively weak due to persistent adaptation. Conversely, as we increase dissimilarity between the adaptation and deviant objects in a parametric way, response recovery should progressively increase to non-adapted levels.

## Methods

### Participants

Twenty-one healthy volunteers (15 females; M age =25.7, SD age = 5.8, range 18 - 40), participated in the fMRI experiment. All participants were right-handed and had normal or corrected-to-normal vision. Written informed consent was obtained from all participants before starting the study. Participants were monetarily compensated for their participation when they completed the experiment. The study was approved by the Ethics Committee of the Faculty of Psychology and Educational Sciences of the University of Coimbra in accordance with the Declaration of Helsinki. We excluded five, four, and two runs (out of ten) for three participants respectively due to excessive head motion, and two runs for another participant due to technical problems in the scanner.

### Procedures

#### fMRI task

Participants completed two fMRI sessions (separated by at least a week), with five runs in each session. We used an event-related design adapted from Experiment 1a, using the same adaptors and deviants. It is important to note that participants were not instructed to perform the one-back task to avoid response confounds; instead, they were asked to passively view the images. Each run started and ended with 16-seconds (s) of a centralized fixation cross. Within each run, there were 50 trials, including 10 of each deviant type (SC, C, D, SD, and I) and 10 of each adaptation length (i.e., five, six, seven, eight, or nine). Each trial had five to nine adaptation objects, followed by two deviants. All adaptor-deviant pairs were presented in each run. Each stimulus (adaptation or deviant) was presented for 500 ms, followed by 500 ms of a blank screen (see Figure 1C and D).

Stimulus delivery and response collection were controlled by “A Simple Framework” (Schwarzbach, 2011) based on the Psychophysics Toolbox on Matlab R2019a (The MathWorks Inc., Natick, MA, USA). Stimuli were presented on an Avotec projector with a refresh rate of 60 Hz and viewed by the participants through a mirror attached to the head coil inside the MR scanner bore. For all experiments, we used an eye tracker to (subjectively) monitor the individual’s attention (and wakefulness) during the task. Although participants were completely naïve, we instructed them to pay attention to the stimuli because the experimenter would ask questions at the end. Subsequently, we administered an informal questionnaire of 21 objects and asked them to identify (by crossing them with a pen) the objects they had seen in the scanner. These data were not analyzed and were only collected to ensure that participants were attentive during the task.

#### MR parameters

MRI data were acquired using a 3T MAGNETOM Trio whole-body MR scanner (Siemens Healthineers, Erlangen, Germany) with a 64-channel head coil at the University of Coimbra (Portugal; BIN - National Brain Imaging Network). Structural MRI data were collected using T1-weighted rapid gradient echo (MPRAGE) sequence (repetition time (TR) = 2000 ms, echo time (TE) = 3.5 ms, slice thickness = 1 mm, flip angle = 7 deg, field of view (FoV) = 256⨯256 mm^2^, matrix size = 256⨯256, bandwidth (BW) = 190 Hz/px, GRAPPA acceleration factor 2). Functional MRI (fMRI) data were acquired using a T2*- weighted gradient echo planar imaging (EPI) sequence (TR =1000 ms, TE = 30 ms, voxel size = 3⨯3⨯3 mm^3^, slice thickness = 3 mm, FoV = 210⨯210 mm^2^, matrix size = 70⨯70, flip angle=68 deg, BW=2164 Hz/px, GRAPPA acceleration factor 2), using a multiband factor of 3 (MB=3). Each image volume consisted of 42 contiguous transverse slices recorded in interleaved ascending slice order oriented parallel to the anterior-posterior commissure plane covering the whole brain. Physiological signals (cardiac and respiratory) were collected.

### Data Preprocessing

#### Anatomical

Both anatomical and functional data were pre-processed using fMRIPrep 20.2.3. This pipeline includes standard preprocessing steps, organized in an easy-usage workflow that ensures robustness independently of the data idiosyncrasies, and has high consistency of results (Esteban et al., 2019). The T1-weighted (T1w) images were corrected for intensity non-uniformity (INU) with N4BiasFieldCorrection (Tustison et al., 2010), distributed with ANTs 2.3.3 (Avants et al., 2008). The T1w-reference was then skull-stripped with a *Nipype* implementation of the ANTS BrainExtraction.sh workflow, using OASIS30ANTs as target template. Brain tissue segmentation of cerebrospinal fluid, white-matter and gray-matter was performed on the brain-extracted T1w using fast (FSL 5.0.9; Zhang et al., 2001). A T1w-reference map was computed after registration of T1w images (after INU-correction) using mri_robust_template (FreeSurfer 6.0.1; Reuter et al., 2010). Brain surfaces were reconstructed using recon-all (FreeSurfer 6.0.1; Dale et al., 1999), and the brain mask estimated previously was refined with a custom variation of the method to reconcile ANTs-derived and FreeSurfer-derived segmentations of the cortical gray-matter of Mindboggle (Klein et al., 2017).

#### Functional

A reference volume and its skull-stripped version were generated using custom methodology from *fMRIPrep*. Susceptibility distortion correction was omitted. The BOLD reference was then co-registered to the T1w reference using bbregister (FreeSurfer) which implements boundary-based registration (Greve and Fischl, 2009). Co-registration was configured with six degrees of freedom. Head-motion parameters with respect to the BOLD reference (transformation matrices, and six corresponding rotation and translation parameters) were estimated before any spatiotemporal filtering using mcflirt (FSL 5.0.9; Jenkinson et al., 2012). BOLD runs were slice-time corrected using 3dTshift from AFNI (Cox and Hyde, 1997). The BOLD time-series (including slice-timing correction when applied) were resampled onto their original, native space by applying the transforms to correct for head-motion. The BOLD time-series was resampled into standard space, generating a *preprocessed BOLD run in MNI152NLin2009cAsym space*. Automatic removal of motion artifacts using independent component analysis (ICA-AROMA; Pruim et al., 2015) was performed on the *preprocessed BOLD on MNI space* time-series after removal of non-steady state volumes and spatial smoothing with an isotropic, Gaussian kernel of 6mm FWHM (full-width half-maximum). Corresponding “non-aggressively” denoised runs were produced after such smoothing. The confound time series derived from head motion estimates and global signals were expanded with the inclusion of temporal derivatives and quadratic terms for each (Satterthwaite et al., 2013).

### Analysis

#### Adaptation analysis

A random-effects approach within the general linear model in SPM (SPM12 - Wellcome Trust Centre for Neuroimaging, London, UK) was used to calculate adaptation effects. There were fourteen regressors: one per adaptor stimulus (from one to nine) and five deviant conditions (including the Identity condition). Although we regressed the five deviants, they were not used in this analysis. We included 24 nuisance regressors: six motion parameters and 18 physiological regressors of cardiac and respiratory cycles. Nuisance regressors were analyzed using the PhysiO Toolbox in SPM (Kasper et al., 2017), including six cardiac, eight respiratory, and four regressors for the interaction between respiratory and cardiac. We modelled the hemodynamic response for each voxel using a general linear model consisting of finite impulse response (FIR) functions. For each regressor of interest, delta functions representing the response following stimulus presentation at each full-volume echo-planar acquisition in a one-second window were fitted to the MR signal. This resulted in an estimate of the response to a single stimulus of each stimulus type, with no assumptions about the shape of the hemodynamic response.

Subsequently, we calculated the areas where the activation elicited by the fourth adaptor was greater than the seventh adaptor (adaptation effect) using an *F*-test for repeated measures - the results were thresholded at p < 0.001 (uncorrected). We decided to contrast these two adaptation object positions to enable an increase in the object-related signal (hence the fourth adaptation stimulus) and, then, compared this signal with one of the alternative adaptation stimuli while maximizing the number of trials that included that length. We opted for the seventh adaptation stimulus instead of the ninth, as only the longest adaptation trials would include this stimulus. Moreover, previous studies have shown that in rapid event-related designs, hemodynamic response peaks around four seconds after the first image presentation (i.e., in our case, four images in a row), followed by a consistent decrease in the signal (see similar results for rapid event-related design in Grill-Spector et al., 2006; Yang et al., 2016).

To ensure the validity of our adaptation results, we assessed the areas that presented a greater BOLD activation in response to a different object compared to a different exemplar of the adapted objects. This was achieved by contrasting deviants (SC, C, D, and SD) against the Identity condition. Although we chose not to include this analysis in our main pipeline to avoid potential circularity issues (Kriegeskorte et al., 2009), results are available in the supplementary materials and, as expected, the areas obtained with this contrast were similar to the areas that exhibited adaptation as measured above.

#### Release from adaptation analysis

We employed a general linear model in SPM using a hemodynamic response function (HRF) with six regressors of interest: one for all adaptors and one for each deviant. Additionally, we included 24 nuisance regressors: six motion parameters and 18 physiological regressors of cardiac and respiratory cycles. Nuisance factors were calculated in the same manner as in the adaptation analysis.

Subsequently, betas from the four deviants were piled into a 4D image in the SC to SD direction. It is important to note that we excluded the Identity condition from our analysis as we wanted to see neural tuning curves as a function of similarity between different objects without the confound of having an Identity condition. Then, using Matlab R2021a (The MathWorks Inc., Natick, MA, USA), we calculated a voxel-wise linear regression inside the areas exhibiting adaptation. In other words, we used the areas that adapted as a mask, and, within these areas, we searched for voxels exhibiting an increase in BOLD response from SC to SD (i.e., according to our hypothesis). This analysis created an R^2^ (i.e., explained variance) map for each participant. Finally, we used the Randomize one sample t-test with 5000 permutations, corrected by threshold-free cluster enhancement (TFCE; FSL package, version 6.04), and controlled for multiple comparisons (*p*FWE-corrected < 0.001). From the areas demonstrating statistically significant R^2^ values, we retained those presenting a positive slope. The regions that did not present a positive slope were mainly parietal lobe, frontal areas, as well as the lateral regions of the ventral stream. We decided not look closely at these regions that did not exhibit positive slope because there is no a priori theoretical hypothesis for inspecting these voxels.

#### Cluster Analysis

We used cluster analysis to assess whether the similarity-driven tuning curves presented the same shape across all regions that showed a release from adaptation. Initially, we estimated the optimal number of clusters using the Calinski-Harabasz index (Calinski and Harabasz, 1974). Then, we used a k-means clustering algorithm to divide the areas into the number of clusters defined by the index according to the shape of their tuning curves.

We extracted the beta activations for each subject and cluster. Subsequently, we Z-transformed the beta activations to compare adaptation and release. Then, for the release analysis, we calculated the slope of the increase from SC to SD for each participant and cluster, and we performed paired t-tests to compare the slopes of clusters 1 and 2. Finally, within each cluster, we conducted paired t-tests to compare the differences between deviants and corrected them for multiple comparisons using Bonferroni. In the cluster graphs, we present the Identity condition beta activation to illustrate that it is lower than the beta activations for the other four deviants.

## Results

We first determined which regions showed a decrease in BOLD response to the adaptation objects. We computed a univariate whole-brain contrast to identify voxels where the fourth adaptation stimulus elicited stronger responses than the seventh – i.e., searching for those areas that showed a decrease in BOLD response due to the repetition of (different exemplars and/or perspectives of) an object type (e.g., a fork; see Figure S3 of the supplementary material for the adaptation curves).

As depicted in Figure 3A, we observe neural adaptation in regions within the ventral occipitotemporal cortex bilaterally, extending from the LOC to the parahippocampal gyrus, and dorsal occipital regions extending to the superior and inferior parietal cortex and the intraparietal sulcus (IPS). There are also more sparse adaptation effects within the superior, middle, and inferior frontal gyri and supplementary motor area (see Figure 3A; blue outline – adaptation effect). To confirm these results, we also contrasted the deviants (SC, C, D, and SD) with the Identity condition (see Figure S2 of the supplementary material), and the areas that adapted were very similar. In subsequent analyses we focused on the adaptation regions obtained with the former method (i.e., the adaptation period; 4^th^ > 7^th^ adaptor).

**Figure 3:**
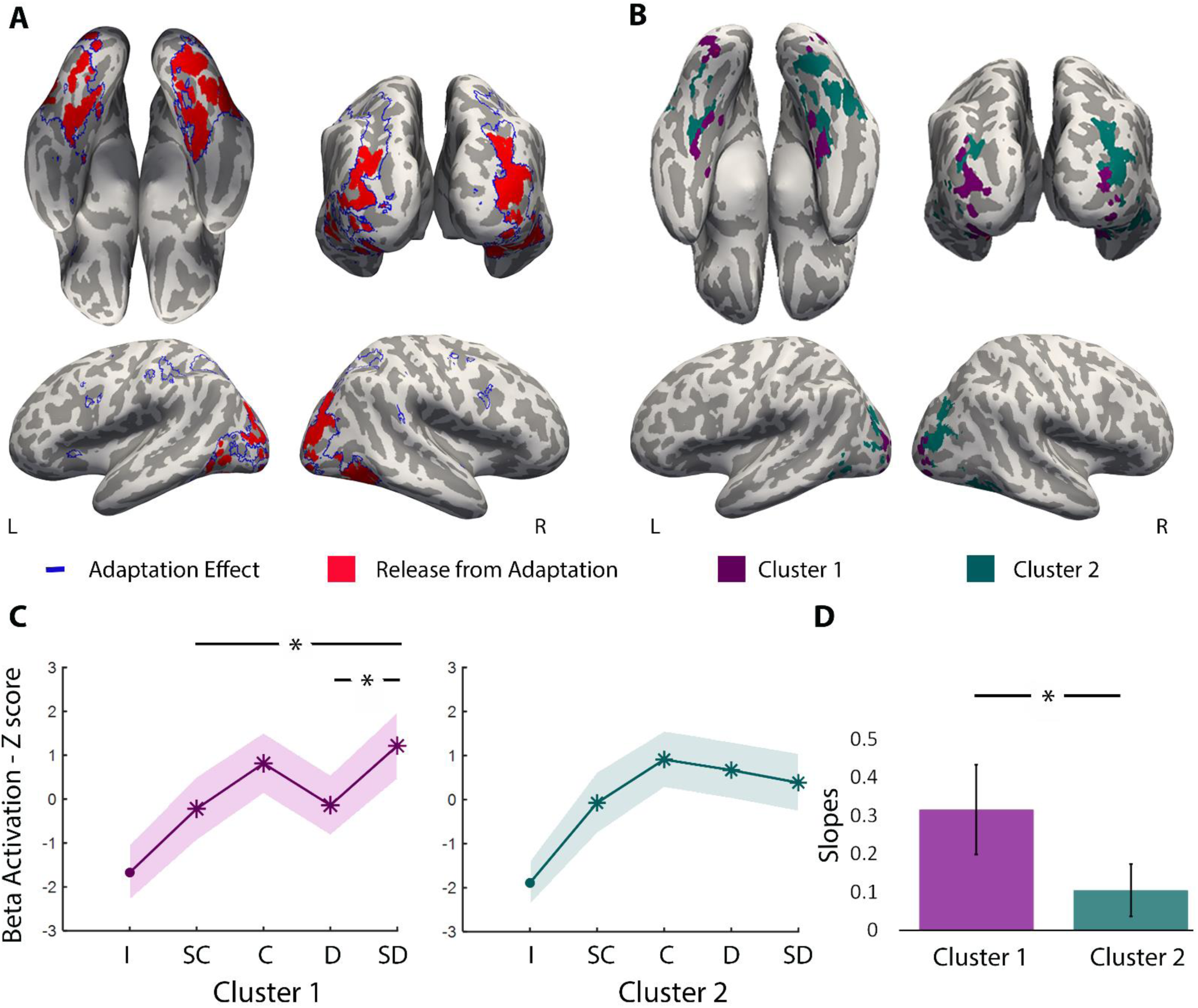
Areas that exhibit adaptation and release from adaptation as a function of similarity between adaptation and deviant objects. **A)** The blue outline encompasses areas that exhibit adaptation (4^th^ > 7^th^ adaptor) at a threshold of p < 0.001 (uncorrected). Those areas were within ventral occipitotemporal cortex (that goes from LOC to parahippocampal gyrus), parietal lobe (inferior and superior) and occipito-parietal cortex, and frontal lobe (superior, middle, and inferior frontal gyrus, and supplementary motor area). In red, we present areas that show release from adaptation as a function of object similarity (*p*FWE-corrected < 0.001). Bilaterally, these include the collateral sulcus and fusiform gyrus (posterior to anterior), the parahippocampal gyrus, the lingual gyrus, the inferior temporal gyrus, the middle temporal gyrus, LOC and the most posterior part of the parietal lobe and occipito-parietal cortex. **B)** Here we show the areas belonging to the two different clusters. Cluster 1 comprises LOC, occipito-parietal cortex, lingual gyrus, parahippocampal, collateral sulcus and the most anterior region of medial fusiform; Cluster 2 comprises parts of the fusiform gyrus, middle temporal gyrus, inferior temporal gyrus, and bilateral occipito-parietal cortex and posterior/caudal IPS. **C)** The graphs represent the release effect in BOLD activation (Z-score and SEM) as a function of the four deviants for the areas of cluster 1 (in purple) and cluster 2 (in green; * p < 0.05 Bonferroni corrected). The graphs were Z-scored to enable comparison between adaptation and release. The Identity (I) condition was never used in our analysis, presented here to show visually that beta activation is below the main conditions in the two clusters (in comparison with the deviants, *p* is consistently below 0.05). **D)** The graph illustrates the differences in slopes between clusters 1 and 2 using a paired t-test (*p = 0.02). SC – Super Close; C – Close; D-Distant; SD – Super Distant

We then tested which of these regions showed graded release from this adaptation as a function of object similarity. To do so, we first computed a voxel-wise linear regression (not including the Identity condition) in search for voxels that showed a linear increase in release from adaptation from the SC to the SD condition. As can be seen in Figure 3A (in red), release from adaptation is present in medial aspects of the ventral temporal cortex along the collateral sulcus, posterior to anterior, from the lingual gyrus to the parahippocampal gyrus. We also observed release from adaptation within the lateral temporal cortex, in the posterior middle temporal gyrus, and finally, in the dorsal occipital and posterior parietal cortical regions. All the release from adaptation effects were bilateral (see Figure 3A; red areas).

Subsequently, we evaluated the characteristics of the neuronal tuning curves in regions showing release from adaptation. We split the data into two clusters according to the Calinski-Harabasz optimal number and applied k-means clustering with two centroids, revealing two distinct similarity-driven tuning curves. Cluster 1 included voxels bilaterally within the medial fusiform, collateral sulcus, parahippocampal gyrus, LOC, and occipito-parietal cortex (see Figure 3B and C in purple). Cluster 2 showed average tuning curves that are more modest in their graded release from adaptation, including voxels in more lateral areas of the fusiform gyrus, occipito-temporal, occipito-parietal cortex, and caudal IPS (see Figure 3B and C in green). Beta values were extracted for each participant and deviant type, and we compared the slopes between clusters and the beta values across deviants for each cluster. We identified that one participant exhibited beta values three standard deviations greater than the group average. Thus, this participant was identified as an outlier, and consequently excluded from these comparisons. However, the results, including this outlier participant, are available in the supplementary material (see Figure S4) and are not generally different from those presented here. In an exploratory analysis, we compared the slope of cluster 1 (M = 0.32; SEM = 0.12) and cluster 2 (M = 0.10; SEM = 0.07) using a paired t-test and found that cluster 1 is statistically steeper than cluster 2 (*t*(19) = 2.49, *p* = 0.02; see Figure 3D). That is, areas in cluster 1 show a progressive release from adaptation as one traverses the different levels of similarity between the adaptation and deviant objects from very similar to very dissimilar. Moreover, Z-score beta activation for cluster 1 indicated statistically lower values for SC (M = −0.22, SEM = 0.70) than SD (M = 1.22; SEM = 0.74; *t*(19) = −3.61, *p* = 0.002; see Figure 3C), and D than SD (M = −0.14; SEM = 0.66; *t*(19) = −2.98, *p* = 0.008). Other comparisons were not statistically significant after Bonferroni correction: SC and C (M = 0.81; SEM = 0.67; *t*(19) = −2.38 *p* = 0.03), SC and D (*t*(19) = −0.17, *p* = 0.87), C and D (*t*(19) = 2.23, *p* = 0.04), C and SD (*t*(19) = −0.95, *p* = 0.36). In cluster 2, there were no statistically significant results after correction for multiple comparisons.

Importantly, the release observed in cluster 1 seems to align more closely with the behavioral data obtained, as it exhibits a steeper slope and a significant difference between SC and SD. Therefore, we focused more on this cluster. In Figure 4, we present the adaptation and release graphs for the areas within cluster 1. Additionally, we include the behavioral results from Experiments 1a and 1b to visually demonstrate the congruency between behavioral and neural data. Interestingly, our results point to an inverse relationship between the behavioral and neural findings. When two objects were more similar (i.e., SC condition), participants tended to be slower and less accurate in detecting the object, and this was reflected in a weaker release effect at the neural level. In contrast, when two objects were more distinct (i.e., SD condition), participants tended to be faster and more accurate behaviorally, and there was a stronger release effect at the neural level.

**Figure 4:**
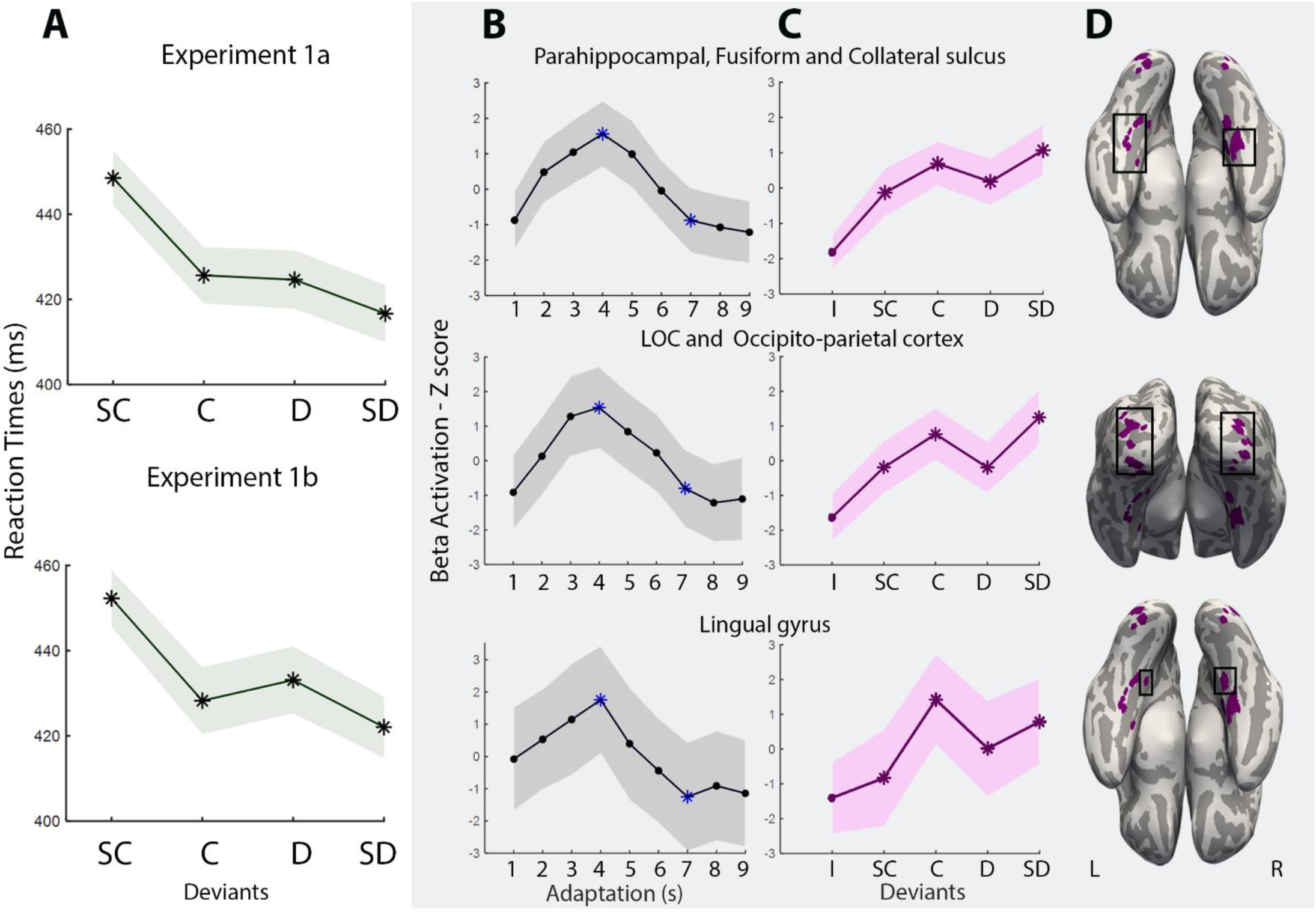
Comparison of the behavioral results and the neural similarity-driven tuning curves for the areas of cluster 1. **A)** The results of behavioral reaction times for Experiment 1 (a and b) - the line represents the average, and the shadow represents the SEM. **B)** Adaptation curves for each brain area, where we present the beta activation Z-scores. We show the BOLD activation for the nine adaptors, and the blue asterisks represent the contrast (4^th^ > 7^th^) to measure adaptation. **C)** For the deviants, we present the beta-activation Z-scores for the five conditions: Identity (I; was never used in our analysis, but is present to show that beta activation is below the main conditions); Super Close (SC); Close (C); Distant(D); and Super Distant (SD). The graphs were Z-scored to enable comparison between adaptation and release. The asterisks represent the conditions we used to calculate behavioral and neural responses. For columns B and C, the line represents the average BOLD amplitude, and the shadow represents the SEM. **D)** Areas of cluster 1.

## Discussion

In Experiment 2, as expected, we observed a gradual reduction in BOLD activation due to the repetitive presentation of an adapting stimulus (different exemplars and/or perspectives of the same object). The bilateral areas demonstrating adaptation effects included the occipito-temporal cortex, parietal lobe, occipito-parietal cortex, frontal areas (superior, middle, and inferior frontal gyri), and supplementary motor area). These areas are known to be preferentially activated by manipulable objects (e.g., Chao et al., 1999; Mahon et al., 2007; Almeida et al., 2013; Bergström et al., 2021). Importantly, we found a release in the BOLD response as a function of object similarity in some areas that presented adaptation. These areas encompassed the fusiform gyrus, collateral sulcus, parahippocampal gyrus, lingual gyrus, inferior temporal gyrus, middle temporal gyrus, LOC, the most posterior part of the parietal lobe, and occipito-parietal cortex (see Figure 3A). Voxels within these areas appear to be tuned to object similarity.

Importantly, these different areas are organized into two distinct clusters based on the tuning curves for the similarity between the adaptation and deviant stimuli. Particularly, the medial fusiform gyrus, collateral sulcus, parahippocampal gyrus, the ventral part of LOC, occipito-parietal cortex, and lingual gyrus formed one cluster (cluster 1; see Figure 4 B, C, and D). This cluster exhibited a more pronounced graded effect of object similarity on the magnitude of the BOLD release, revealing a clear distinction between SC and SD, and seems to align more closely with our behavioral data in Experiment 1 (a and b). In contrast, ventral temporal areas situated more posterior and lateral than those in cluster 1, as well as dorsal occipital regions that are more superior and anterior than those in cluster 1, belonged to another cluster as a function of their tuning function (cluster 2). These areas displayed a more modest graded tuning function in response to progressive manipulations of object similarity – visually, there appears to be a distinction between the SC condition and all other deviants in this cluster (see Figure 3B and C). These differences may relate to the processing preferences of the clustered areas and, potentially, the processing level at which these areas operate.

## General Discussion

Our ability to recognize and discriminate one object from other objects in the environment is crucial for completing basic daily tasks. To understand this ability, we may need to unravel how object knowledge is organized in the brain – and, specifically, we may need to focus on finer-grain representations and test whether, at this level of representation, similarity is a major organizing principle of information in the brain (Op De Beeck et al., 2008; Grill-Spector and Weiner, 2014). Here, we show that object similarity impacts object within-category representations – these representations may be anchored in object similarity in a relatively graded way, perhaps in the same way as sensory-motor regions code for low-level perceptual similarity. This observation seems to be particularly true for the medial fusiform gyrus and collateral sulcus, the parahippocampal gyrus, the lingual gyrus, and LOC – i.e., major ventral stream regions important for object recognition and processing (Grill-Spector and Weiner, 2014) – as well as occipito-parietal cortex.

Behaviorally, we demonstrated that participants faced greater difficulty in detecting an object change when the adaptation object and the deviant object were more similar. This implies that detecting an object becomes more challenging in the presence of similar objects, suggesting higher confusability between objects with greater object similarity. Analogous results have been consistently observed in numerical processing. Specifically, performance in a number decision task – e.g., deciding if a target numerosity set was different than the preceding one – varied as a function of the distance between the two sets. Parenthetically, it has been shown that number similarity, described by Weber’s law, governs neural tuning curves in the intraparietal sulcus (Piazza et al., 2004; Harvey et al., 2013; Kersey and Cantlon, 2017).

Notably, our results show that neural responses in regions of cluster 1 (i.e., parahippocampal and medial fusiform gyri, collateral sulcus, lingual gyri, LOC, and occipito-parietal cortex) seem to be tuned to the full spectrum of object similarity. Moreover, the shape of their release tuning curve aligns with that obtained in our behavioral experiments, suggesting some involvement in our ability to detect and recognize a different object – i.e., the increasing similarity between two deviant and adaptation objects led to progressively slower reaction times and progressively weaker recovery of the BOLD response. Areas of this cluster have been shown to be involved in aspects of high-level manipulable object processing. Specifically, they play a crucial role in processing object-specific visual information, such as information on shape, material, and surface properties (Grill-Spector and Malach, 2001; Hayworth and Biederman, 2006; Cant and Goodale, 2007; Cant et al., 2009; Pyles and Grossman, 2009; Gallivan et al., 2014; Freud et al., 2017b; Almeida et al., 2023a). Additionally, the medial fusiform gyrus has been associated with motor-relevant object information and functional grasping (Mahon et al., 2007; Valyear and Culham, 2010; Chen et al., 2016; Lee et al., 2019; Knights et al., 2022; Almeida et al., 2023a; Mahon and Almeida, 2024), whereas the parahippocampal and fusiform gyri have an important role in processing semantic knowledge and object function (Chao et al., 1999; Wheatley et al., 2005; Aminoff et al., 2013; Mandera et al., 2017; Kleineberg et al., 2018; Almeida et al., 2023a). In fact, the medial fusiform gyrus might work as a central area for processing information about manipulable objects and shows privileged functional connectivity with the parietal lobe, premotor cortex, and lateral occipitotemporal cortex (Mahon et al., 2007, 2013; Almeida et al., 2013; Stevens et al., 2015; Garcea et al., 2016; Kristensen et al., 2016; Chen et al., 2017; Amaral et al., 2021; Walbrin and Almeida, 2021 see also Mahon and Caramazza, 2011, for a functional description of this connectivity see Mahon and Almeida, 2024). Finally, occipito-parietal regions within this cluster are involved in object-specific 3D processing (Georgieva et al., 2008; Kilintari et al., 2011; Almeida et al., 2014, 2023a; Freud et al., 2016) and object grasping (Jeannerod et al., 1994; Davare et al., 2011; Monaco et al., 2011; Almeida et al., 2023a).

A second cluster of areas shows a more modest graded release from adaptation and seems to have only a very crude representation of similarity. This cluster includes areas more posterior and superior to those in cluster 1, such as the posterior occipital-temporal and superior occipito-parietal cortex, which are also involved in processing shape, texture, and motor functions (Peuskens et al., 2004; Wurm and Lingnau, 2015; Freud et al., 2017a; Almeida et al., 2023a). However, these regions potentially encode less complex and lower-level features, which might not allow for fine-grain representations necessary for fully differentiating objects or for encoding a comprehensive object space. Interestingly, the ventral temporal regions associated with clusters 1 and 2 seem to follow an organization from anterior and medial to posterior and lateral: regions in cluster 1 are more medial and anterior, whereas regions in cluster 2 are more posterior and lateral. Perhaps this posterior-to-anterior difference reflects the putative hierarchical processing of objects and how this processing relates to increasingly fine-grained processing of object features and object knowledge. On the one hand, lower levels of analysis, whereby very crude processing of object knowledge leads to rather rudimental similarity effects (i.e., making it harder to see graded similarity differences), are present in more posterior and lateral regions. On the other hand, as the level of analysis increases in complexity and we move more anteriorly and medially, the processing of object features becomes more fine-grained, and concomitant graded effects of feature-based object similarity become more apparent. A key aspect of this paper involves the use of a release from adaptation technique. Many current studies on similarity have used multivariate approaches across stimuli to explore the role of similarity in object processing (e.g., Kriegeskorte et al., 2008; Carlson et al., 2014; Contini et al., 2020; Fernandino et al., 2022). However, we firmly believe that this technique is the most suitable design to measure similarity-based tuning curves and show the graded nature of similarity as a principle in organizing finer-grained representations - as potentially seen in other types of information processing (Piazza et al., 2004; Fabbri et al., 2010; Planton et al., 2021). The nature of this technique allows us to see individual effects of progressive increase in object similarity in the processing of objects. Moreover, by using release from adaptation, we can conduct behavioral and neural experiments with a similar structure, facilitating the comparison of reaction times and BOLD recovery. Additionally, adaptation is likely a mechanism that aids in conserving energy and enhances the sensitivity of the neural population to new stimuli (Grill-Spector et al., 2006), making release from adaptation probably more sensitive to fine-grained distinctions. Studies using adaptation have significantly contributed to our understanding of how fine-grained information is represented in the human brain. One of the most prominent examples is the numerosity studies (Piazza et al., 2004, 2007; Cantlon et al., 2006), which paved the way for later ground-breaking discoveries, such as identifying the numerosity maps in the intraparietal sulcus (Harvey et al., 2013). Note that although adaptation and release from adaptation techniques are still widely used nowadays, some authors have raised questions about the difficulties in interpreting these effects (Larsson et al., 2016).

Importantly, in our behavioral and neural tuning curves, particularly in areas within cluster 1, there is a clear distinction between SC and SD within the category of manipulable objects. However, the results are less conclusive at intermediate levels (i.e., C and D). Unfortunately, due to the limited number of data points (only four), we cannot make more definite assumptions about the shape of the tuning curve, but our data seems to suggest that the complete tuning curve may not be entirely linear, possibly because the similarity values in these conditions are not far enough apart to result in appreciable behavioral and neural differences (see Table S1 of the supplementary material). One potential solution to unravel these intermediate levels is to conduct the same experiments with more deviants to obtain enough data points for curve fitting. Nevertheless, the results of both experiments present evidence of release from adaptation consistent with strong cognitive and neural tuning functions for fine-grained object similarity. Furthermore, our findings indicate that this coding reflects a relatively continuous graded object similarity space, which is analogous, perhaps, to face space and face adaptation results (Jiang et al., 2009).

Finally, we purportedly employed a relatively broad measure of similarity that combined all object features irrespective of their content. As such, we cannot say, with certainty, what types of content are driving similarity effects in behavioral responses or the graded similarity releases in the different brain areas. Although we acknowledge the involvement of these areas in visual processing (Grill-Spector and Malach, 2001; Hayworth and Biederman, 2006; Cant and Goodale, 2007; Cant et al., 2009; Pyles and Grossman, 2009; Gallivan et al., 2014; Freud et al., 2017b), we consider it unlikely that we are only assessing this aspect, especially, because several studies have shown that these same areas might be central for processing various others types of knowledge (Chao et al., 1999; Wheatley et al., 2005; Mahon et al., 2007; Valyear and Culham, 2010; Aminoff et al., 2013; Chen et al., 2016; Mandera et al., 2017; Lee et al., 2019; Knights et al., 2022; Almeida et al., 2023a). Additionally, our adaptation paradigm presented different exemplars and perspectives of the same object, leading us to believe that participants were adapting to the object itself and not specifically to low- to mid-level features such as orientation. Thus, in the same vein, the recovery of the BOLD response should depend not only on these low- to mid-level features but also on other significant object properties (visual or not), such as texture, material, action knowledge, or other conceptual information.

Overall, our results demonstrate that information about manipulable objects is represented in such a way as to maintain real-object similarity in a relatively graded way, both cognitively and neurally. This supports the hypothesis that object similarity reflects real-world proximity and that finer-grained representations within a category retain a rather detailed and graded map of similarity. This representation likely plays a central role in how the brain process and recognize everyday objects.

## Acknowledgements

We would like to thank Fernanda Ponce and Angelika Lingnau for their important input in the early stage of this study.

## Supplementary Material

**Supplementary Table S1:** Pairs used in the Behavioral Experiments, in brackets there is the cosine similarity between adaptor and each deviant.

|  | Adaptors | Deviants |  |  |  |
| --- | --- | --- | --- | --- | --- |
|  |  | SC | C | D | SD |
| Experiment 1a | Hoe | shovel<br>(0.76) | plunger<br>(0.41) | door handle<br>(0.10) | cork<br>(0.01) |
|  | Drill | drill bit<br>(0.74) | hole punch<br>(0.46) | paint brush<br>(0.10) | hanger<br>(0.03) |
|  | Scissors | knife<br>(0.75) | grater<br>(0.47) | sponge<br>(0.09) | match<br>(0.06) |
|  | Fork | spoon<br>(0.71) | peeler<br>(0.41) | match<br>(0.10) | cork<br>(0.06) |
|  | Pliers | swiss knife<br>(0.67) | screw<br>(0.42) | board eraser<br>(0.10) | napkin<br>(0.06) |
|  | Bucket | bottle<br>(0.64) | bowl<br>(0.42) | nutcracker<br>(0.09) | horn<br>(0.03) |
|  | Glass | bottle<br>(0.90) | cup<br>(0.54) | weights<br>(0.10) | horn<br>(0.03) |
|  | Bottle Cap | cork<br>(0.76) | bottle opener<br>(0.43) | weights<br>(0.10) | candle holder<br>(0.04) |
|  | Hand-Blender | grater<br>(0.78) | peeler<br>(0.42) | mop<br>(0.10) | candle holder<br>(0.06) |
|  | Stapler | paper clip<br>(0.75) | rubber stamp<br>(0.41) | spinning top<br>(0.10) | plunger<br>(0.06) |
| Experiment 1b | Shovel | hoe<br>(0.76) | mop<br>(0.6) | bottle opener<br>(0.27) | sponge<br>(0.08) |
|  | Knife | scissors<br>(0.75) | wooden spoon<br>(0.40) | pastry bag<br>(0.21) | pencil eraser<br>(0.08) |
|  | Bottle | glass<br>(0.90) | bowl<br>(0.51) | citrus squeezer<br>(0.25) | horn<br>(0.02) |
|  | Broom | mop<br>(0.76) | plunger<br>(0.54) | pencil<br>(0.20) | door handle<br>(0.06) |
|  | Spoon | fork<br>(0.71) | peeler<br>(0.42) | clothespin<br>(0.19) | shaver<br>(0.07) |
|  | Grater | hand blender<br>(0.78) | pliers<br>(0.40) | hammer<br>(0.21) | board eraser<br>(0.07) |
|  | Cork | bottle cap<br>(0.76) | bottle opener<br>(0.31) | rolling pin<br>(0.10) | candle holder<br>(0.01) |
|  | Swiss Knife | knife<br>(0.71) | grater<br>(0.42) | plunger<br>(0.20) | napkin<br>(0.06) |
|  | Nail clipper | scissors<br>(0.64) | shaver<br>(0.38) | hammer<br>(0.17) | paddle<br>(0.04) |
|  | Paper clip | stapler<br>(0.75) | screw<br>(0.41) | clothespin<br>(0.18) | paddle<br>(0.03) |

**Supplementary Figure S1:**
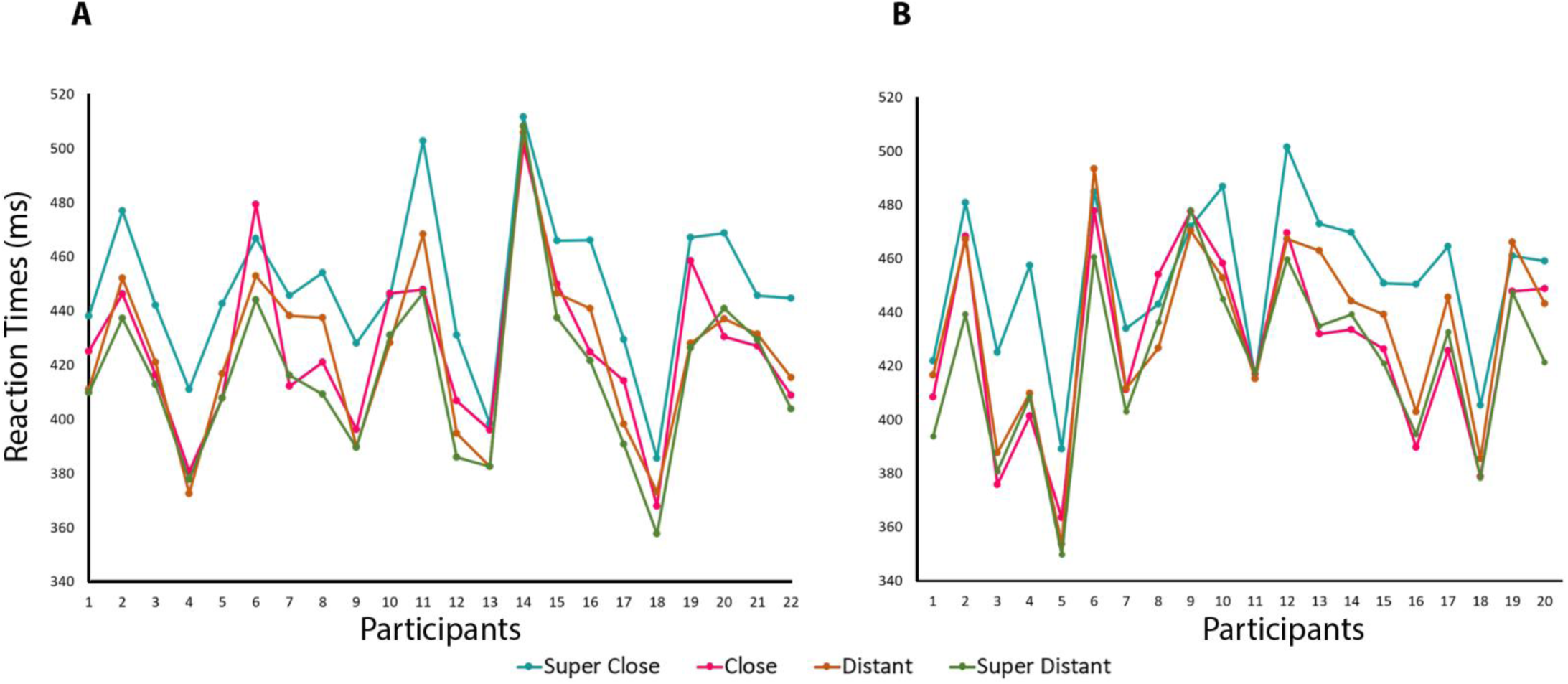
Reaction Times (in ms) per deviant type and participant in A) Experiment 1a and B) Experiment 1b. The blue line represents the Super Close condition, the pink line represents the Close condition, orange represents Distant, and green represents Super Distant conditions.

**Supplementary Figure S2:**
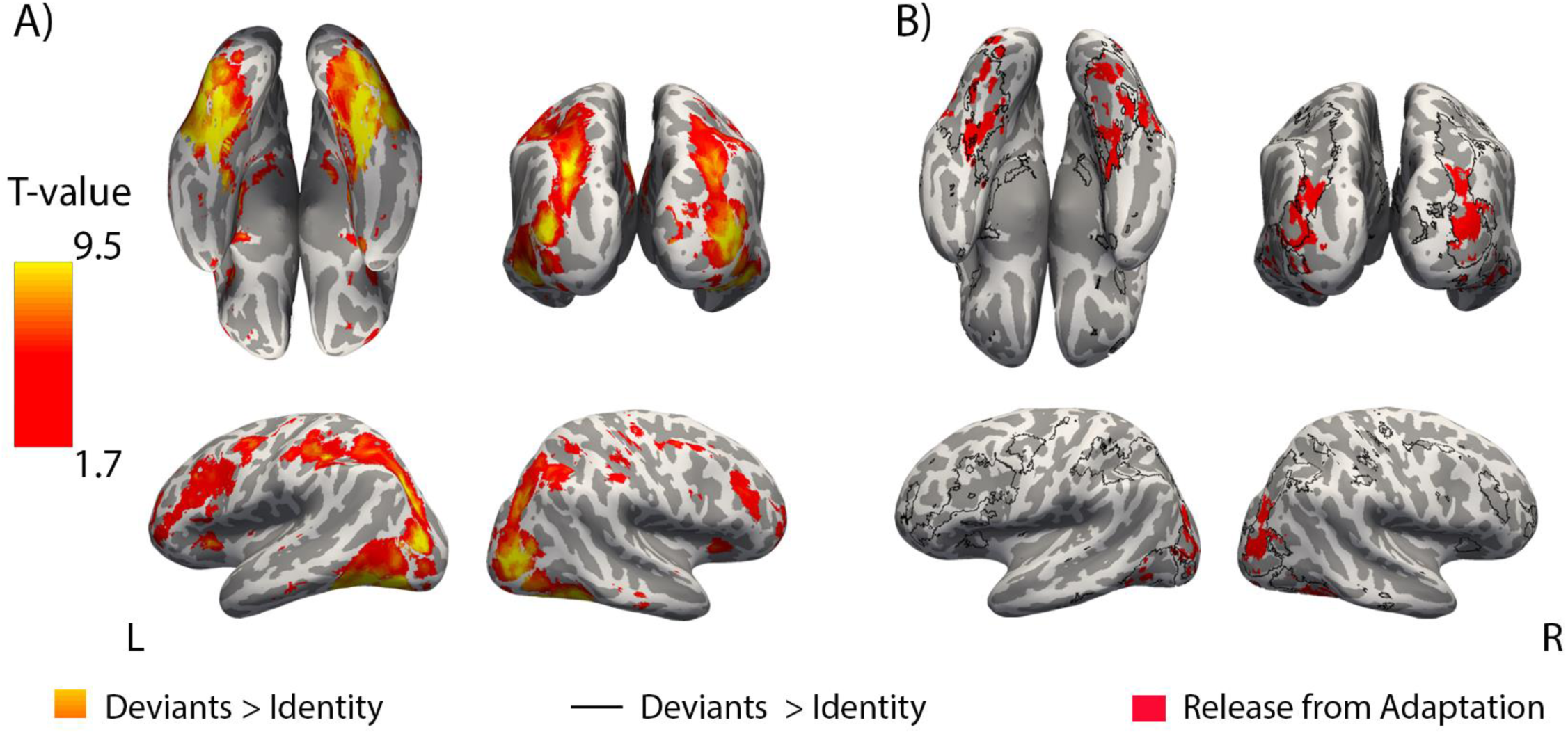
Areas presenting a difference between the deviants and the Identity condition and their relationship with the areas that present release from adaptation. **A)** A paired t-test was conducted between the four deviants (i.e., SC, C, D, and SD) and the Identity condition, *p* < 0.05 (uncorrected). **B)** The figure illustrates the overlap between the contrast Deviants > Identity (in black outline) and the areas that have a release from adaptation (in red).

**Supplementary Figure S3:**
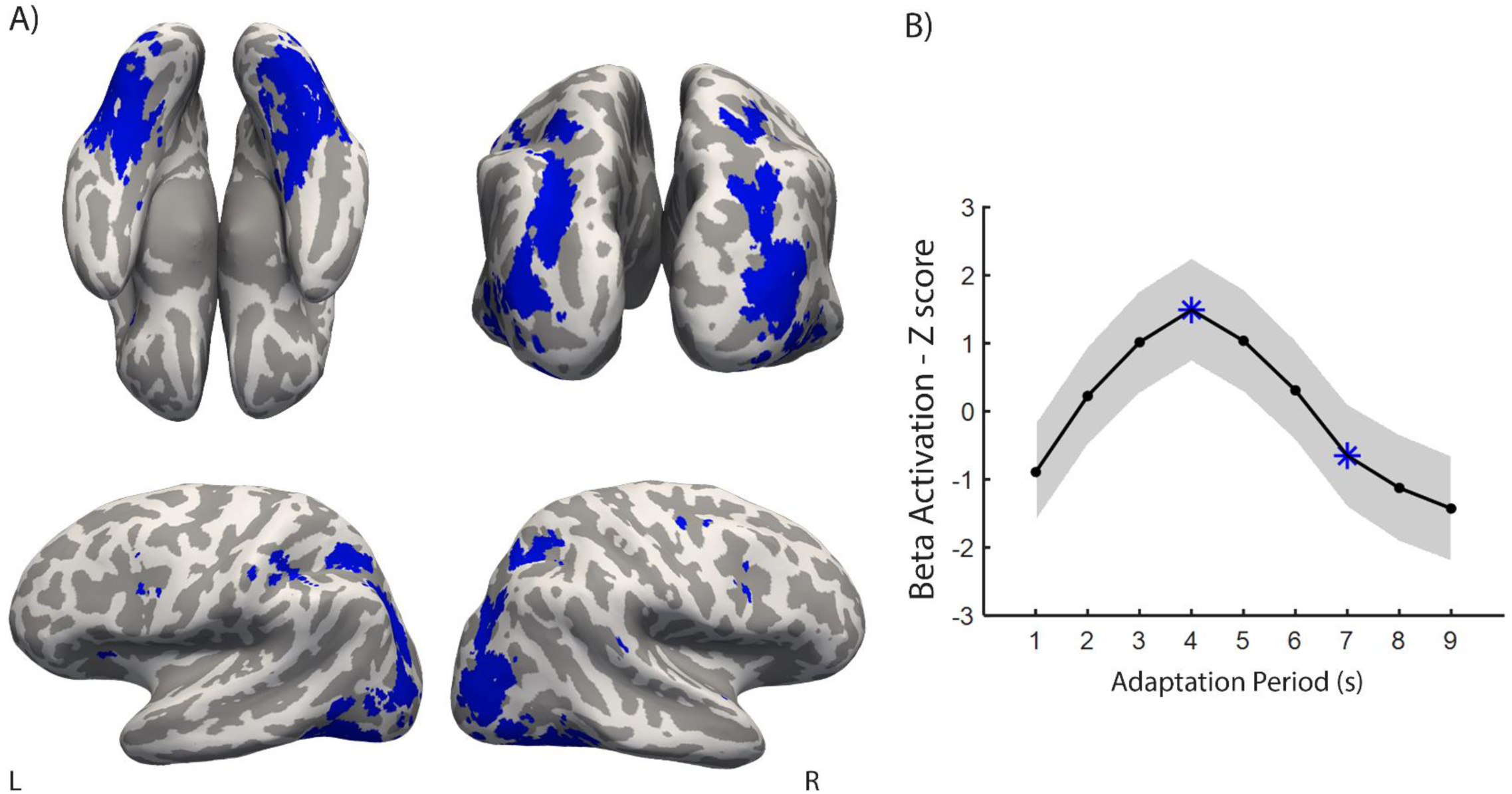
fMRI-adaptation. A) Areas that presented adaptation defined as 4^th^ > 7^th^ adaptor under p < 0.001 (uncorrected). These areas are presented in the paper in Figure 3A, blue outline. B) The line graph represents the BOLD adaptation effect (Z-score and SEM) as a function of the nine adaptors. The blue asterisks show the two moments that we used to define adaptation.

We evaluated the characteristics of the two neuronal tuning curves in the regions that showed release from adaptation (see Figure S4). Without excluding the outlier participant, we compared the slope of cluster 1 (M = 0.24; SEM = 0.13) and cluster 2 (M = 0.07; SEM = 0.07) using a paired t-test, but it did not reach statistical significance (*t*(20) = 1.86, *p* = 0.08). However, using a non-parametric test – the Wilcoxon Signed-Rank test, we found that cluster 1 is statistically steeper than cluster 2 (*Z* = 1.96, *p* = 0.05; see Figure S4D).

When conducting paired t-tests to compare beta values across deviants for each cluster, we did not find statistical significance after correction for multiple comparisons. Yet, using a more lenient approach with paired t-tests one-tailed, we found statistically significant results. In cluster 1, beta activation values are statistically lower for SC (M = −0.16, SEM = 0.79) than C (M = 0.96, SEM = 0.86; *t*(20) = −2.70, *p* = 0.007, one-tailed), and SD (M = 0.99; SEM = 0.72; *t*(20) =-2.81, *p* = 0.005, one-tailed; see Figure S4C). Other comparisons were not statistically significant after Bonferroni correction, such as SC and D (M = −0.04; SEM = 0.78; *t*(20) = −0.26, *p* = 0.40, one-tailed), C and D (*t*(20) = 2.53, *p* = 0.01, one-tailed), C and SD (*t*(20) = −0.05, *p* = 0.48, one-tailed), and D and SD (*t*(20) = −2.19, *p* = 0.02, one-tailed). Using a similar approach, in cluster 2, the only statistically significant result was the difference between SC (M = −0.05; SEM = 0.64) and C (M = 0.94; SEM = 0.63), showing that SC is lower than C (*t*(20) = −2.74, *p* = 0.006, one-tailed; see Figure S4C). Other comparisons were not statistically significant after Bonferroni correction: SC and D (M = 0.70; SEM = 0.62; *t*(20) = −2.39, *p* = 0.01, one-tailed), SC and SD (M = 0.28, SEM = 0.58; *t*(20) = −1.30, *p* = 0.10, one-tailed), C and D (*t*(20) = 0.93, *p* = 0.18, one-tailed), C and SD (*t*(20) = 2.21, *p* = 0.02, one-tailed), and D and SD (*t*(20) = 1.36, *p* = 0.09, one-tailed).

**Supplementary Figure S4:**
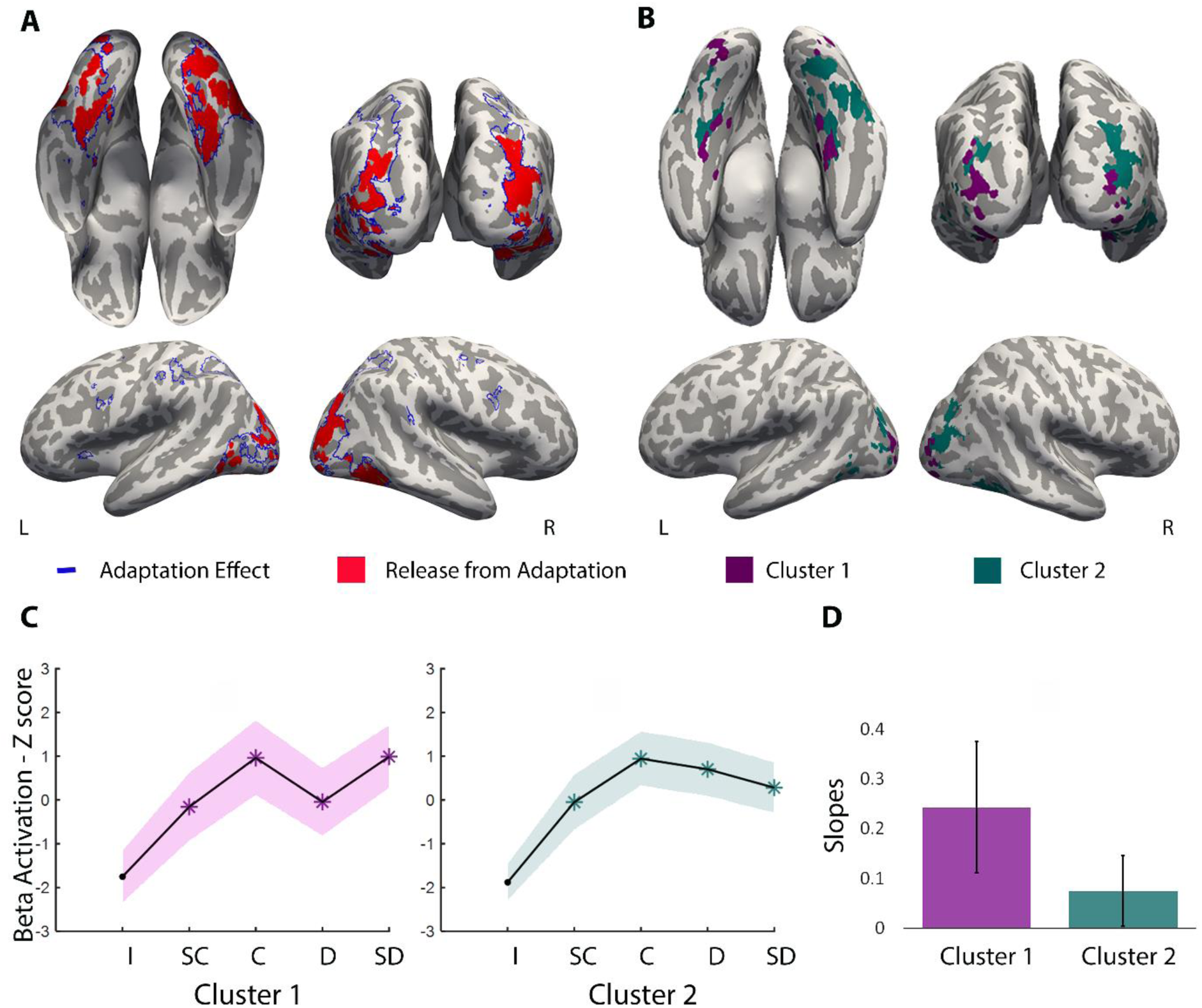
Areas that exhibit adaptation and release from adaptation as a function of similarity between adaptation and deviant objects without excluding the outlier. **A)** The blue outline encompasses areas that exhibit adaptation (4^th^ > 7^th^ adaptor) at a threshold of p < 0.001 (uncorrected). Those areas were within ventral occipitotemporal cortex (that goes from LOC to parahippocampal gyrus), parietal lobe (inferior and superior) and occipito-parietal cortex, and frontal lobe (superior, middle, and inferior frontal gyrus, and supplementary motor area). In red, we present areas that show release from adaptation as a function of object similarity (*p*FWE-corrected < 0.001). Bilaterally, these include the collateral sulcus and fusiform gyrus (posterior to anterior), the parahippocampal gyrus, the lingual gyrus, the inferior temporal gyrus, the middle temporal gyrus, LOC and the most posterior part of the parietal lobe and occipito-parietal cortex. **B)** Here we show the areas belonging to the two different clusters. Cluster 1 comprises LOC, occipito-parietal cortex, lingual gyrus, parahippocampal, collateral sulcus and the most anterior region of medial fusiform; Cluster 2 comprises parts of the fusiform gyrus, middle temporal gyrus, inferior temporal gyrus, and bilateral occipito-parietal cortex and posterior/caudal IPS. **C)** The graphs represent the release effect in BOLD activation (Z-score and SEM) as a function of the four deviants for the areas of cluster 1 (in purple) and cluster 2 (in green). Identity (I) condition was never used in our analysis, presented here to show that beta activation is below the main conditions in the two clusters. **D)** The graph illustrates the differences in slopes between clusters 1 and 2.

